# Generation of CD34^+^CD43^+^ hematopoietic progenitors to induce thymocytes from human pluripotent stem cells

**DOI:** 10.1101/2021.09.02.458682

**Authors:** Léa Flippe, Anne Gaignerie, Céline Sérazin, Olivier Baron, Xavier Saulquin, Ignacio Anegon, Laurent David, Carole Guillonneau

## Abstract

Immunotherapy using primary T cells has revolutionized medical care in some pathologies in recent years but limitations associated to challenging cell genome edition, insufficient cell number production, the use of only autologous cells and lack of product standardization have limited its uses in the clinic. The alternative use of T cells generated in vitro from human pluripotent stem cells (hPSCs) offers great advantages by providing a self-renewing source of T cells that can be readily genetically modified and facilitate the use of standardized universal off-the-shelf allogeneic cell products and rapid clinic access. However, despite their potential, the feasibility and functionality of T-cells differentiated from hPSCs needs better comprehension before moving to the clinic. In this study, we generated human induced pluripotent stem cells from T-cells (T-iPSCs) allowing preservation of already recombined TCR, with the same properties as human embryonic stem cells (hESCs). Based on these cells, we differentiated with high efficiency hematopoietic progenitor stem cells (HPSCs), capable of self-renewal and differentiation into any cell blood type, and then DN3a thymic progenitors from several T-iPSC lines. To better comprehend differentiation, we analyzed the transcriptomic profiles of the different cell types and demonstrated that HPSCs differentiated from hiPSCs had a very similar profile to cord blood hematopoietic stem cells (HSCs). Furthermore, differentiated T-cell progenitors had a similar profile to thymocytes at the DN3a stage of thymic lymphopoiesis. Therefore, with this approach, we were able to regenerate precursors of therapeutic human T cells to potentially treat a wide number of diseases.

## INTRODUCTION

Immunotherapy with T effector (Teff) or T regulatory (Treg) cells offers new possibilities to treat diseases like cancer or autoimmune diseases and transplantation rejection in need for new therapeutics (Bézie et al., 2017, 2019a; Themeli et al., 2015). Next-generation therapeutics using Teff or Treg cells have even started to emerge thanks to genetic engineering and the possibility to provide an antigen specificity to the genetically modified T cells by expressing TCRs or chimeric antigen receptors (CARs) to improve the outcome (Bézie et al., 2019b; Flippe et al., 2019; Nianias & Themeli, 2019). However, the clinical use of Teffs or Tregs has only been done from autologous T cells and this raises problems to produce them in the required time window. This autologous source can also limit the number of T cells that can be produced in some instances. The development of “off-the-shelf” allogeneic T cells is awaited as the next generation T cells that will overcome these obstacles. An approach to generate “off-the-shelf” T lymphocytes is to develop an ex vivo platform that will mimic lymphopoiesis starting with stem cells and generate T cells.

Lymphopoiesis, a process resulting from the generation of progenitors from the bone marrow and their maturation in the thymus, is difficult to study in human and recapitulate *in vitro*. To date, the best option is a scRNAseq trajectory reconstruction providing the possibility to map T cells development (Park et al., 2020). Although hematopoietic stem cells (HSCs) can differentiate into T cells (Awong et al., 2011; De Smedt et al., 2004; La Motte-Mohs et al., 2005; Schmitt & Zúñiga-Pflücker, 2002), they cannot be amplified beforehand or edited efficiently. Pluripotent stem cells are an alternative source with great potential for the production of T cells. In particular, hiPSCs reprogrammed from somatic cells, are more accessible than hESCs and offer the opportunity to develop therapies based on antigen specificity.

Reprogramming from T cells has been shown to generate T-iPSCs with pre-existing V(D)J rearrangements at the T cell receptor loci; in turn, using T-iPSCs could facilitate the generation of fully competent T cells. Indeed, the antigenic specificity can be retained during reprogramming. It has been shown that the reprogramming of MART1-specific human T cells generates T-iPSCs that can subsequently differentiate into functional T cell clones with the same specificity as the original cell and functional *in vitro* and *in vivo* (Kawamoto et al., 2018a; Maeda et al., 2016). Similarly, it has also been shown in mice that adoptive transfer of CD4^+^ Tregs derived from autoantigen-specific T-iPSCs significantly reduced the CD8^+^/CD4^+^ ratio in the pancreas of diabetic mice (M. Haque et al., 2019). However, differentiation of T cells from hPSCs remains difficult, therefore a better understanding of the differentiation process is required.

To address this issue, we generated T-iPSCs and subjected those cells to T cell differentiation and compared them with several hPSC lines. Next, we analyzed hematopoietic stem progenitor cells (HSPCs) differentiated from hPSCs vs CD34^+^ cord blood HSCs and revealed that the CD34^+^CD43^+^ subset is the most closely related to CD34^+^ cord blood HSCs. We then subjected HPSCs from hPSCs and CD34^+^ cord blood cells to T cell differentiation with primary thymocytes as a control to clearly pinpoint stage match differentiated cells with in vivo situation and demonstrated efficient differentiation up to the DN3a T cell development stage. Therefore, our work lays down the basis for studying the establishment of TCR expression during T cell differentiation in humans, as well as a potential method to produce large number of thymocytes in which engineered TCRs or CARs can be expressed.

## RESULTS

### Efficient establishment of hPSCs clones from blood T cells

We first reprogrammed T-cells from PBMCs from healthy volunteers to obtain T-iPSCs with rearranged TCR regions (Kawamoto et al., 2018b; Maeda et al., 2016). Prior to induction of reprogramming with Sendai OSKM vectors (**Figure 1A**), we activated whole PBMCs for 4 days with αCD3 and αCD28 mAbs and recombinant IL-2, since somatic cell reprogramming is more favorable in cells with an active cell cycle (Plath & Lowry, 2011), and obtained 80% TCR^+^CD3^+^ cells (**Figure 1B**). Two weeks after induction, clones were picked, amplified and after 10 passages (2,5 months) T-iPSCs were validated. All tested clones lost Sendai vectors used for the induction of reprogramming factors, and displayed NANOG, SOX2 and OCT4 expression levels similar to the one of hESCs (H9) (**Figure 1C**). We also performed SNP analysis to confirm cell identity and infer copy number variations. We showed that the reprogrammed T cells harbored the same karyotype than the parental cells (**Figure 1D**). Furthermore, while PBMCs and fibroblasts did not show modification in the allele frequency at the q11.2 region of the chromosome 14, corresponding to the TCR rearrangement zone, reprogrammed T05 and T04 T-cell clones showed a clear modification in this region, indicating a rearranged TCR (**Figure 1E**). Finally, to functionally test the T-iPSCs, we performed a trilineage assay, which consisted in inducing germ layer engagement from hiPSCs. Engagement towards mesoderm, endoderm and ectoderm was evaluated by 3’seq-RNA Profiling (3’SRP) (Charpentier et al., 2021) (**Figure 1F**). This assay confirmed that the reprogrammed T cells could differentiate into mesoderm, endoderm and ectoderm.

**Figure 1:**
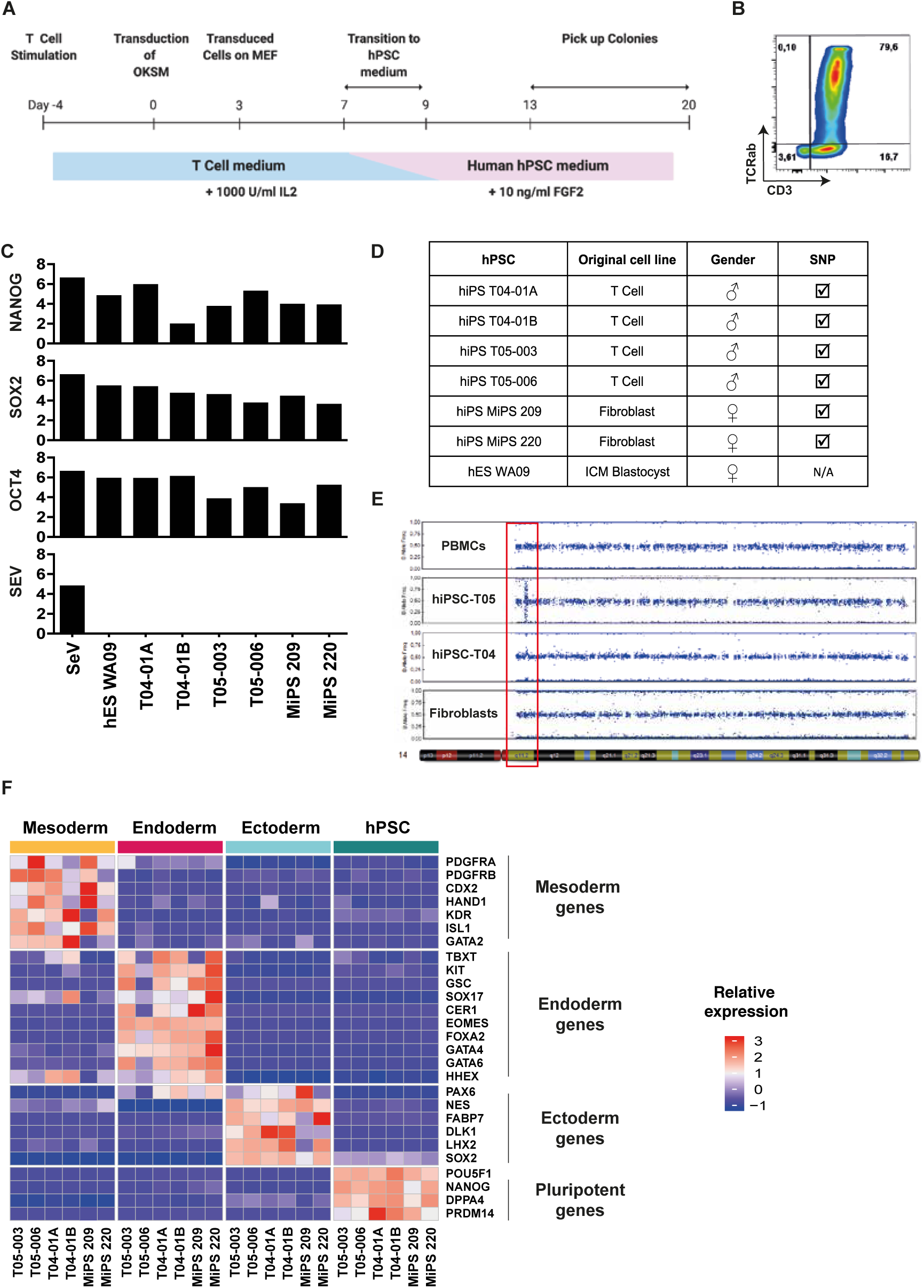
Generation of Human iPSCs from T cells. **A**, Schematic view of the reprogramming protocols. 10^5^ activated T-cells were seeded at day 0, then transduced with 3 Sendai viruses expressing a polycistronic KLF4/OCT4/SOX2, MYC and KLF4 at a ratio of 6:6:3,6 respectively. Cells were split on feeders at day 3 and placed in the hPSCs media at day 9. Colonies started to appear around day 13 and were picked on feeders. **B**, Representative dot plot of CD3 and TCRαβ on T-cells after 4 days of αCD3 and αCD28 mAb activation. **C**, qPCR measurement of Oct4, Nanog, Sox2 and Sendai virus (SEV) expression in indicated cell lines, one positive control (SeV) and one negative control (hES WA09). **D**, Summary of original cell line, gender and SNP profile of indicated cells lines. A tick mark means identical to parental cells. **E**, SNP analysis showing the allelic frequency on chromosome 14 of PBMCs, hiPSC-T05, hiPSC-T04 and hiPSC-MiPS. The red box shows the q11.2 zone representing the TCR rearrangement area. **F**, Indicated cell lines have been differentiated in the three germ layer (mesoderm, endoderm, ectoderm) and have been analyzed by RNseq. Selected gene expression representative of each germ layer were plotted as a heatmap.

In conclusion, we were able to efficiently reprogram PSCs from T cells, therefore generating hiPSC with rearranged TCRs.

### Functionally competent hematopoietic progenitor cells differentiation from reprogrammed T cells

We then subjected hPSCs derived from fibroblast or from T cells to HPSCs differentiation to better understand their respective potential and included hESCs as controls. The differentiation started by mesoderm induction, followed by hemogenic endothelium specification to finally yield CD34^+^CD43^+^ HPSCs at day 9 (**Figure 2A**) (Flippe et al., 2020). Overall, all pluripotent cell lines differentiated efficiently, yielding in average 37% of CD34^+^ cells, although we observed a more efficient generation of CD34^+^ cells using the T04.01B clone vs. the other clones (**Supplementary figure 1A**). For all tested cell lines, the percentage of CD34^+^CD43^+^ population within the CD34^+^ cells evolved drastically between day 7 (less than 1%) and day 9 (around 13%) (**Figure 2C**). Among CD34^+^ cells, CD45^+^ cells increased from 40% at day 7 to 80% at day 9 (**Figure 2B-C**). The clone T04.01B was particularly efficient at differentiating, but since all the other T-cell-derived iPSC clones behaved similarly to hESC and fibroblast-derived iPSCs, we concluded that it was most likely due to a clonal effect (**Supplementary figure 1A-B**).

**Figure 2:**
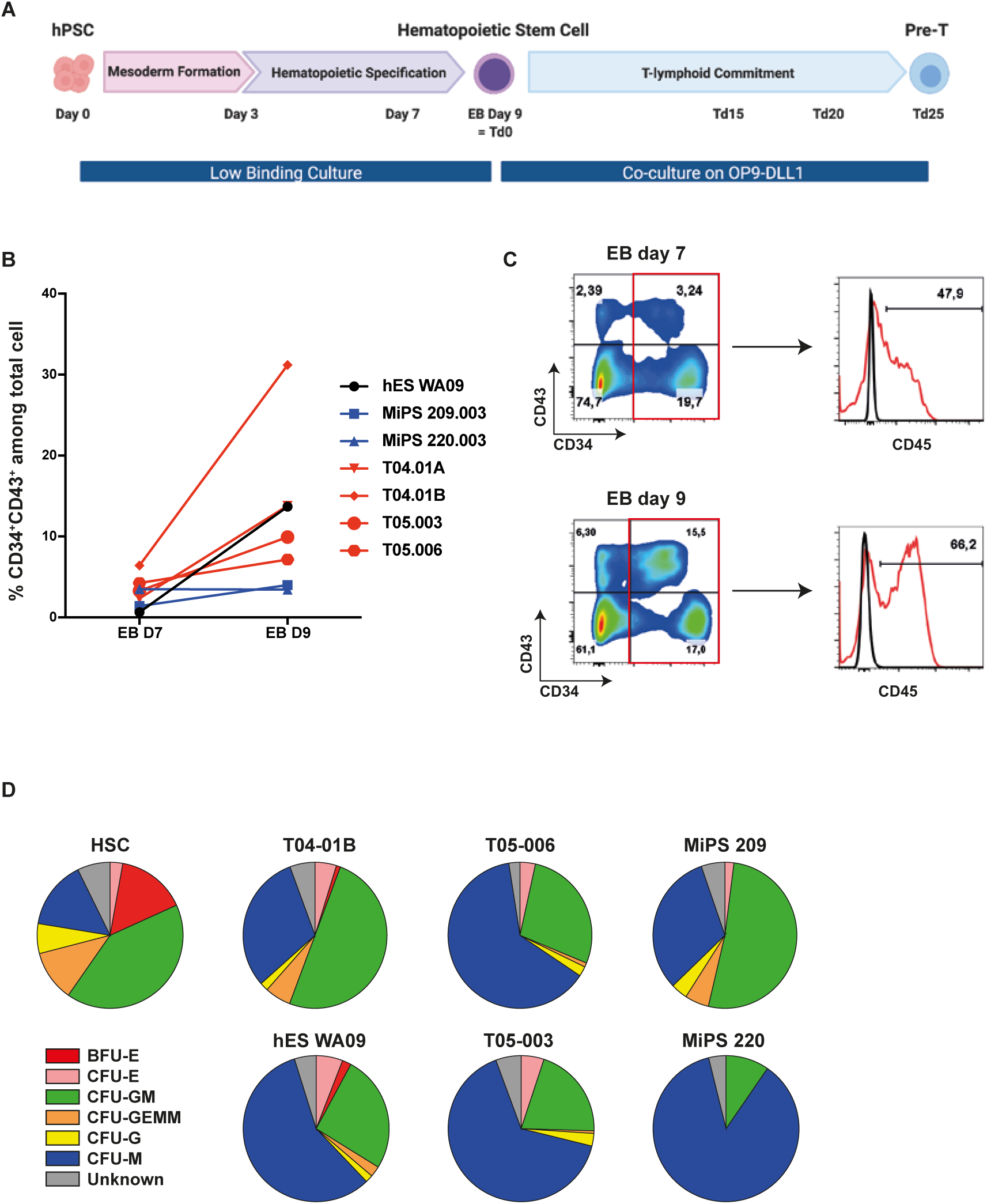
Multipotent HPSCs were derived from hPSCs. **A**, Schematic representation of the hematopoietic stem cell differentiation protocol. Between day 0 and day 9, EBs are treated to induce firstly mesoderm formation then to induce specification towards the hematopoietic lineage. After 9 days, the EBs were dissociated and the cells were co-cultured on OP9-DLL1 or DLL4 cells in specific medium to induce T-lymphoid commitment. **B**, Flow-cytometry analysis of differentiating cells at D7 and D9 in EB culture from hPSC. Representative dot plot of CD34 and CD43 co-staining on living cells is shown. The expression of CD45 in CD34^+^ cells is shown. Red line represents cells stained with a fluorescent antibody and black line represents unstained cells. **C**, Evolution of the percentage of total cells expressing CD34 and CD43 between day 7 and day 9 and expression of CD45. **D**, Summary of visual microscopic counting of CFU assay blood colonies for the indicated cell lines.

To functionally assess CD34^+^ HPSCs, we subjected them to a hematopoiesis assay on methyl-cellulose. Cord-blood HSC CD34^+^ cells subjected to this protocol yielded visible colonies of BFU-E (Burst forming unit-erythroid), CFU-E (Colony forming unit-erythroid), CFU-GM (Colony forming unit-granulocyte, macrophage), CFU-G (Colony forming unit-granulocyte, erythrocyte, macrophage, megakaryocyte), CFU-G (Colony forming unit-granulocyte) and CFU-M (Colony forming unit-macrophage) that can be numerated (**Figure 2D and Supplementary Figure 2A**). After counting the colonies, cells were harvested and expression of erythroid marker (CD235a) and monocyte/macrophage markers (CD14, CD15) were analyzed by flow cytometry (**Supplementary figure 2C**) (Bissels et al., n.d.; Lochem et al., 2004). Starting from similar numbers, we showed that CD34^+^ cells derived from differentiation self-renewed less efficiently than cord blood HSCs (**Supplementary figure 2B**). Nevertheless, the 6 IPS clones tested were able to generate all types of blood colonies, albeit not in the exact same proportions than cord blood HSCs, as shown by morphological numeration (**Figure 2D**) or flow cytometry analysis (**Supplementary figure 2D**). Altogether, this demonstrated that the generated CD34^+^ HPSCs were functional.

### Transcriptomic profiling demonstrates that CD34^+^CD43^+^ derived from hPSC are the most similar to HSCs

To further characterize CD34^+^ HPSCs derived from PSCs, we performed 3’SRP RNAseq on sorted CD34^+^ and CD43^-^ cells from embryoid bodies (EBs) at Day 7, CD34^-^CD43^+^, CD34^-^CD43^-^, CD34^+^CD43^+^, and CD34^+^CD43^-^ cells from EBs at Day 9 (**Supplementary figure 3**) and compared to cord blood HSCs. Hierarchical clustering defined three groups of cells: (1) D7 CD34^+^, D9 CD34^+^ and D9 CD34^+^CD43^-^ EBs; (2) D2 and D5 EBs; and (3) D9 CD34^+^CD43^+^ EBs and cord blood HSCs (**Figure 3A**). We observed that D9 CD34^+^CD43^+^ EBs clustered with cord blood HSCs to form the group of HPSCs. Pearson correlation analysis further confirmed the proximity of D9 CD34^+^CD43^+^ EBs and cord blood HSCs. Surprisingly, D7 CD34^+^ and D9 CD34^+^ EBs showed a high degree of transcriptional similarity, despite relative differences in their potential to become T cells progenitor *in vitro*. Functional enrichment analysis revealed that genes significantly upregulated in EBs D9 CD34^+^CD43^+^ cells were involved in pathways like hematopoietic stem cell and progenitor differentiation, activation of HOX gene during differentiation, NOTCH signaling and Megakaryocyte/Lymphocyte/Leukocyte differentiation (**Figure 3B**). In comparison, genes overexpressed in D9 CD34^+^CD43^-^ EBs were linked to regulation of endothelial cell differentiation, endothelium development, positive regulation of Notch signaling pathway and stem cell development/commitment/differentiation (**Figure 3B**). This global analysis demonstrated that the CD34^+^CD43^+^ subset differentiated from hiPSC was the sub-population more closely related to cord blood HSCs, while in contrast the CD34^+^CD43^-^ subset differentiated from hiPSC was potentially still at the hemogenic-endothelium transition (Vodyanik et al., 2006).

**Figure 3:**
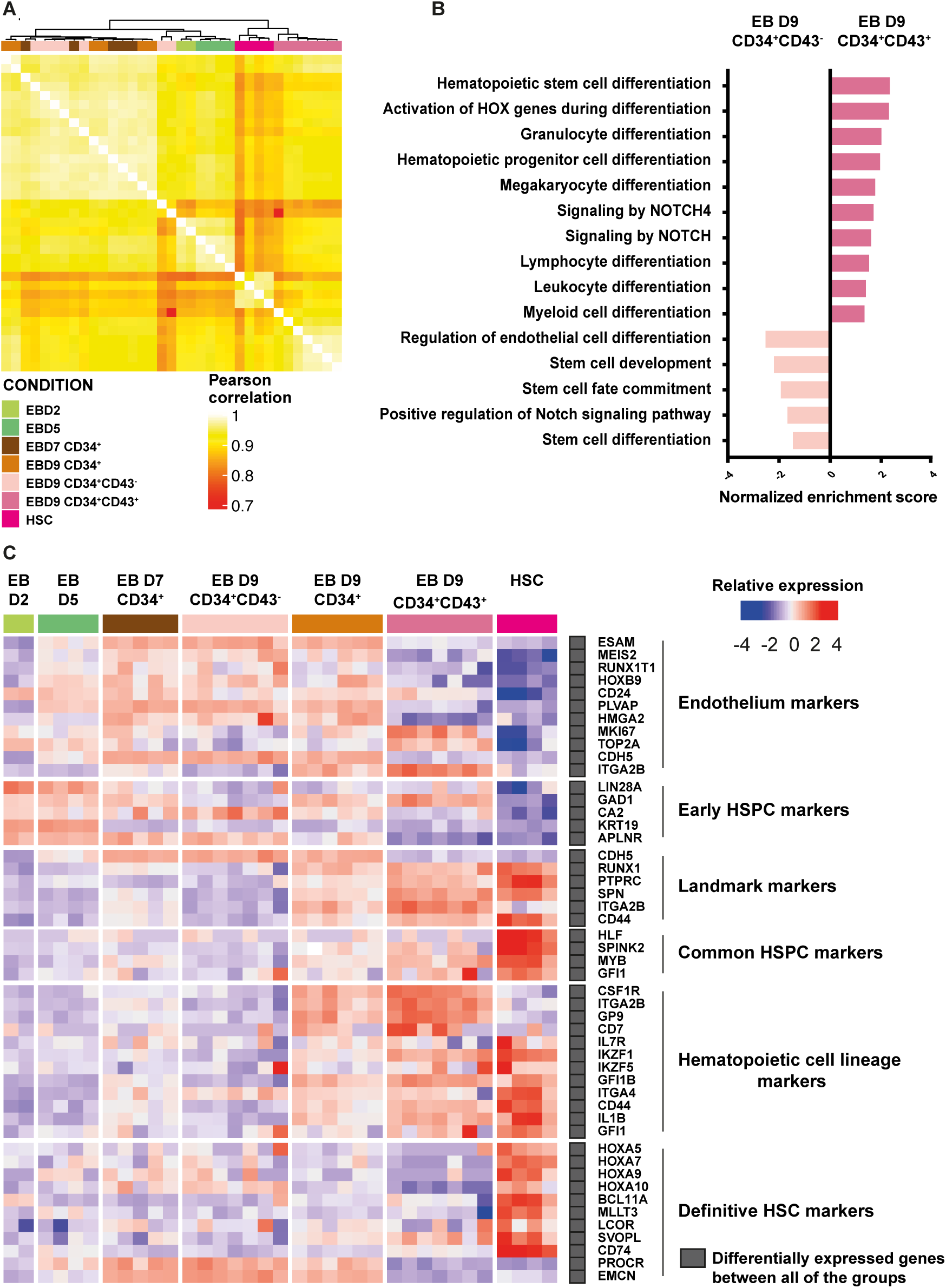
CD34^+^CD43^+^ derived from hPSC are closely related to cord blood HSCs. **A**, Heatmap of Pearson correlation coefficients of HSC and hPSCs differentiated form hPSCs. **B**, Normalized enrichment score of biological pathways up-regulated or down regulated in EB D9 CD34^+^CD43^+^ compare to HSC. All represented pathways have a adjust p-value < 0,05. **C**, Gene expression heatmaps of selected markers characterizing hematopoietic differentiation are shown for digital gene expression sequencing (DGE-seq) for this study.

To further compare HPSCs obtained from iPSCs and cord blood, we analyzed the expression profile of significantly differentially expressed genes subsets associated with the progression of hematopoiesis (**Figure 3C**). We observed that endothelial and early HSPC markers such as ESAM, MEIS2, HOXB9 or CD24 were upregulated in D2, D5, D7 CD34^+^, D9 CD34^+^CD43^-^ and D9 CD34^+^ EBs, while they were downregulated in D9 CD34^+^CD43^+^ EBs and HSCs. Interestingly, MEIS2, a gene involved in the endothelium to hematopoiesis transition (EHT) (M. Wang et al., 2018), was overexpressed by CD34^+^ cells at D7 and CD34^+^CD43^-^ cells at D9 and downregulated by CD34^+^CD43^+^ cells and cord blood HSCs, suggesting that the latter have undergone their EHT. On the contrary, overexpression of well-known markers of HSCs, such as RUNX1, PTPRC (or CD45), CD44, MYB and IKAROS Family Zinc Finger IKZF1 (IKAROS) and IKZF5 (PEGASUS) was observed in D9 CD43^+^CD34^+^ EBs and HSCs, while downregulated in D2, D5, D7 CD34^+^, D9 CD34^+^CD43^-^ EBs. Differentially expressed genes between cells showed hierarchy of acquisition of HPSCs signature. Finally, it was noted that definitive HSC markers such as HOXA (5,9,10) (Sugimura et al., 2017; Vo et al., 2018) and MLLT3 (Calvanese et al., 2019), that have been highlighted for their involvement in self-renewal and engraftment capacity, were upregulated in cord blood HSCs but not expressed in D9 CD34^+^CD43^+^ EBs (**Figure 3C**). Altogether, the transcriptomic signature showed that D9 CD34^+^CD43^+^ EBs were closely related to functional CD34^+^ cord blood HSCs and thus represent the most promising population to support lymphopoiesis.

### Efficient lymphopoiesis induction up to DN3a T cell developmental stage of HPSCs

We then subjected HPSCs and CD34^+^ cord blood HSCs to T cell differentiation using DLL1 or DLL4-overexpressing OP9 cells co-culture. Of note, we recently defined that OP9-DLL1 and OP9-DLL4 were equivalently potent to induce T cell lineage (Flippe et al., 2020). Briefly, D9 EBs were dissociated and seeded on DLL1/4-OP9 cells in a medium supplemented with SCF, FLT3l and IL-7 (Td0) and analyzed after 15 (Td15), 20 (Td20) and 25 days (Td25) (**Figure 2A**). The same treatment was applied to the CD34^+^ cord blood HSCs. After 20 days of co-culture, the expression of the early marker of lymphopoiesis CD7 averaged 68,4% for HSC-Td20 and 70,8% for hPSC-Td20 (**Figure 4A**). In addition, about 9% of CD7^+^ cells expressed the CD4 and CD8 marker in HSC-Td20 versus 3,1% in hPSC-Td20. HSC or hPSC, induced in lymphopoiesis for 20 or 25 days, did not express CD3 and TCRαβ, indicating that the differentiated cells obtained were progenitors.

**Figure 4:**
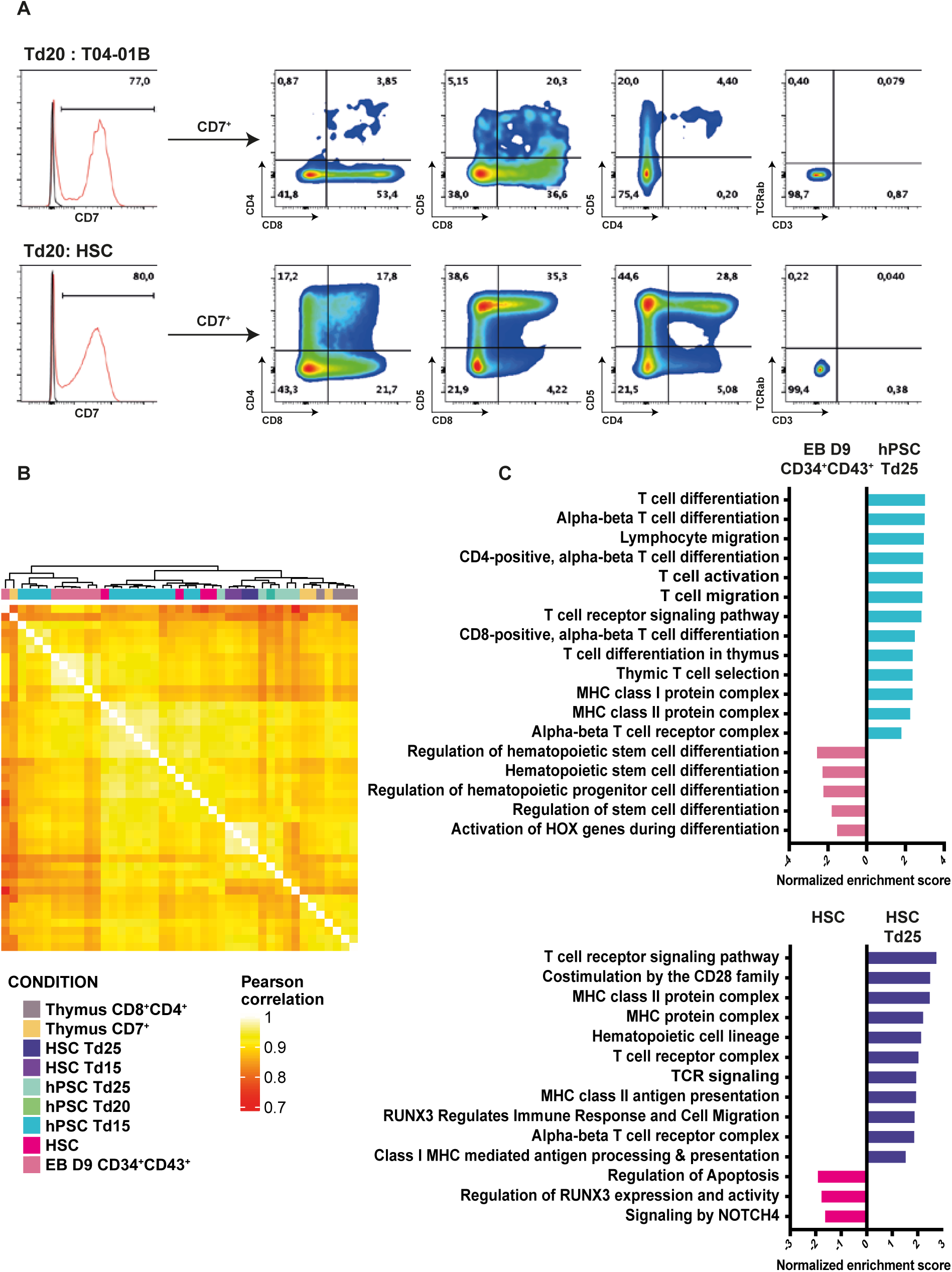
hPSCs-Td25 are closed to CD8^+^CD4^+^ thymocytes. **A**, Representative histogram of CD7 expression in living cells (excluding OP9) at day 29. Red line represents cells stained with a fluorescent antibody and black line represents unstained cells. The expression of CD8, CD4, CD3 and TCRαβamong CD7^+^ cells is shown in dot plot. **B**, Heatmap of Pearson correlation coefficients of thymus cells and hPSCs differentiated form hPSCs. **C**, Normalized enrichment score of biological pathways up-regulated or down regulated in EB D9 CD34^+^CD43^+^ compare to hPSC-Td25 (up) and in HSC compare to HSC-Td26 (down). All represented pathways have a adjust p-value < 0,05.

To further understand the identity of cells differentiated from hPSC or HSCs, we compared their transcriptomic signatures to natural thymocytes. For this, thymocytes were extracted from human thymuses and sorted on CD7 or CD4 and CD8 co-expression (**Supplementary figure 4**). Hierarchical clustering defined four groups of cells: (1) D9 CD34^+^CD43^+^ EBs and Td15, (2) cord blood HSC and Td15, (3) HSC-Td15, HSC-Td25, Td20 and Td25, and (4) CD7^+^ and CD8^+^CD4^+^ thymocytes (**Figure 4B**). Pearson correlation analysis further confirmed a clear distinction between the EB D9 CD34^+^CD43^+^ and the HSC group. HSC were closer to the iPSC-Td15 cells. The iPSC-Td20, iPSC-Td25, HSC-Td15 and HSC-Td25 group was closer to thymocytes. Notably, upon differentiation, analysis highlighted the upregulation of specific pathways related to T cell biogenesis: “alpha-beta T cell differentiation, T cell receptor signaling, thymic T cell selection or MHC class I/II protein complex” and downregulation of pathways such as “hematopoietic stem cell differentiation or regulation of stem cell differentiation” (**Figure 4C**). Similar enrichment was found for HSC and hPSCs differentiation. These results confirmed the commitment of the cells to the T cell lineage.

To further understand the developmental stage achieved by in vitro differentiation of PSCs towards thymic lymphopoiesis, we then analyzed stage-associated makers (**Figure 5**). T cell progression through lymphopoiesis involves progressive phases of lineage specification, characterized by the acquisition of a T cell-specific transcriptional program. The different development stages are characterized as follows: the early intrathymic precursor ‘ETP’ stage, the immediately pre-commitment DN2a stage (DN=double negative for mature T-cell coreceptors CD4 and CD8), the newly committed DN2b stage, the DN3a stage that is the first major period of TCR gene rearrangement, the DN3b and immature single-positive stages when cells proliferate vigorously following the first successful TCR locus gene rearrangement, and finally the DP (CD4, CD8 double positive) stage when the second TCR locus is rearranged and cells successfully express for the first time their final TCR recognition specificity. The DP stage is the progenitor stage that give rise to all effector subsets of T cells using the αβ form of the TCR (Rothenberg et al., 2016). We then analyzed genes well known to define the different developmental stages of T cells in the thymus (Rothenberg et al., 2008; Yui & Rothenberg, 2014; Zhou et al., 2019). Three profiles emerged from this analysis: (1) D9 CD34^+^CD43^+^ EBs and CD34^+^ cord blood HSCs, which overexpressed ETP and DN2a stage-related specific genes and downregulated other stage-related genes, (2) hPSC/HSC Td20, Td25 and thymic cells, which on the contrary downregulated ETP and DN2a stage-related genes and upregulated DN2b, DN3a, DN3b stage-related genes, and (3) hPSCs-Td15 which have a particular intermediate profile (**Figure 5**). We noticed that LMO2, GATA2 and MPO genes involved in the ETP and DN2a stages of differentiation (Buono et al., 2016; Canté-Barrett et al., 2017; Desanti et al., 2011) were upregulated in hiPSC-Td15, while they were downregulated in hiPSC-Td25 and HSC-Td25, as well as in CD7^+^ and CD8^+^CD4^+^ thymocytes. (**Figure 5**). On the other hand, genes such as RUNX3, BCL11B and LEF1 (Garcia-Perez et al., 2020) were found to be upregulated in hiPSC-Td25 and HSC-Td25 and marked the transition to more advanced stages (**Figure 5**). Moreover, we noticed that they also expressed the BCL11B and LCK genes, which are essential for the passage from DN2 to DN3. Interestingly, we observed that hPSC/HSC Td20 and Td25 cells expressed RAG1/RAG2, as well as the 3 chains of CD3 (D/E/G) and PTCRA which are involved in the rearrangement of the TCR complex (Dik et al., 2005) (**Figure 5**). In addition, we could observe that differentiating cells such as CD8^+^CD4^+^ thymocytes expressed DN3b and DP stage-related genes such as TCF7, ZAP70, IKZF1 (IKAROS), IKZF3 (AIOLOS) or ICOS (Dong et al., 2001; Georgopoulos, 2017; Georgopoulos et al., 1997; H. Wang et al., 2010), showing their engagement in the T lineage. (**Figure 5**). Since hPSC/HSC-Td20 and Td25 cells did not express the TCRαβ/CD3 complex protein at the cell surface, that the stage of differentiation of hiPSC-Td25 and HSC-Td25 cells was DN3a (Outters et al., 2015).

**Figure 5:**
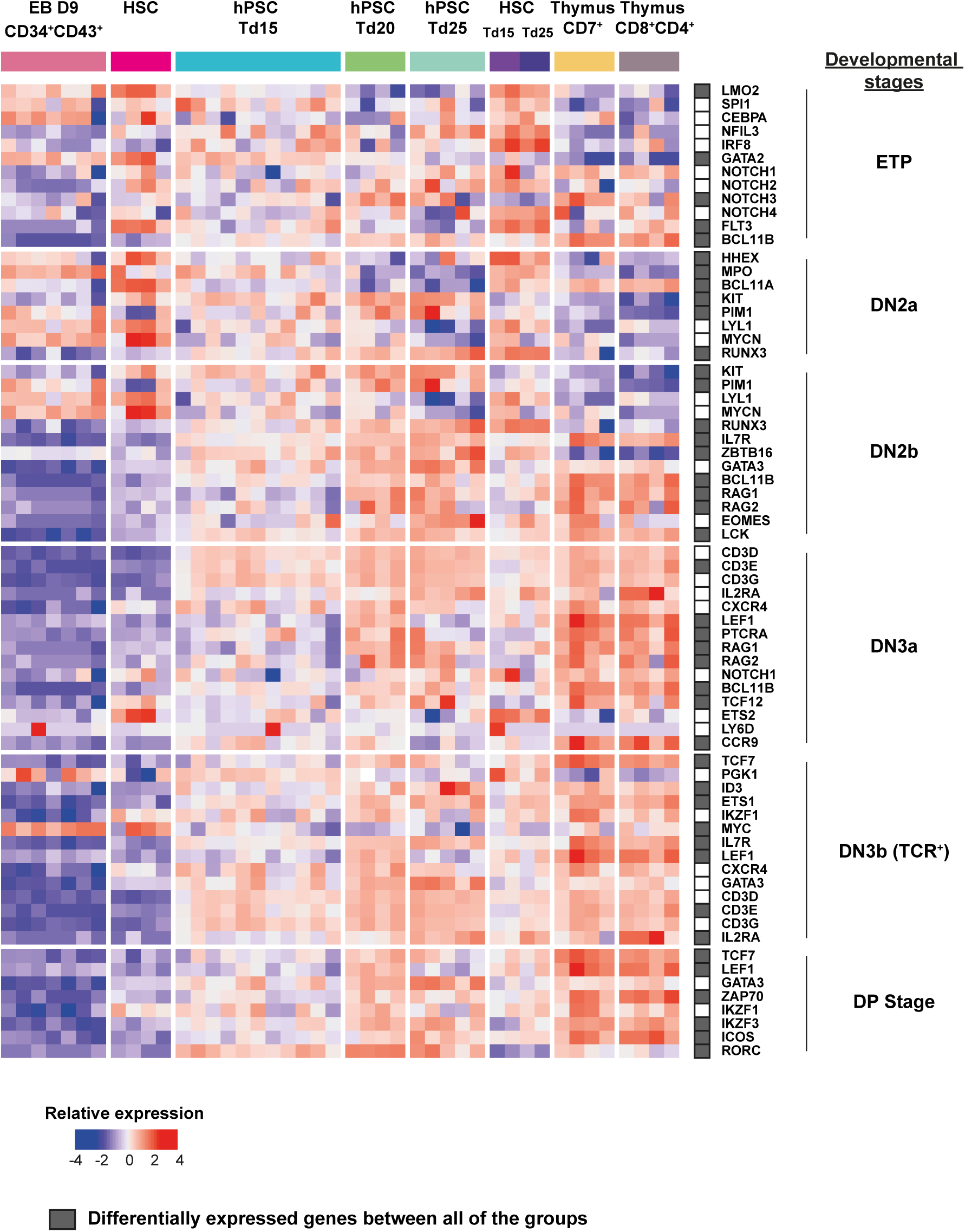
T-cell progenitor derived from hPSC are stuck at the DN3a thymocyte stage. Gene expression heatmaps of selected markers associated in T-cells development and function are shown defined by DGE-seq.

## DISCUSSION

In this study, we successfully and reproducibly generated T cell progenitors at the DN3a stage from several hESC and hiPSC lines. Two differentiation steps were necessary, the first one was the generation of HPSCs from hPSCs through embryonic corpus formation and the second step was the differentiation of HPSCs into T cells through co-culture on OP9-DL1. The comparison of hPSCs-HPSCs with primary cord blood cells confirmed that hPSCs-HPSCs were similar to HSCs. Similarly, we confirmed that hPSC-Td25s were similar to primary thymocytes. Indeed, they recapitulated the transcriptome, protein markers and differentiation potentials of primary cells. For this study, we generated T-iPSCs which were similar to hESCs and other hiPSC lines.

We showed that T-iPSCs could differentiate similarly to hiPSCs reprogrammed from other cell types. Furthermore, it has been demonstrated that T cell reprogramming can increase the lifespan of cells because newly produced cells have longer telomeres, which represents a significant therapeutic advantage (Nishimura et al., 2013). To efficiently direct the differentiation of hiPSCs to the hematopoietic lineage, we used our optimized feeder-free protocol (Flippe et al., 2020). After 9 days of differentiation, we obtained CD34^+^CD45^+^ cells regardless of the hiPSCs or hESCs lines used. Discrepancies between HPSCs and CD34^+^ cord blood have been reported in particular on their multipotency and their ability to engraft in animals (Paes et al., 2017). This can be explained by the fact that HPSCs differentiated from hPSCs recapitulate primitive rather than definitive hematopoiesis (Demirci et al., 2020). Therefore, our first endeavor was to assess HSPC from iPSC vs cord blood. Functional analysis with CFU assay showed similar potential: all CD34^+^, regardless of the source, were able to multiply and differentiate in the whole repertoire of blood colonies. This suggested that the differentiated cells could be fully capable of differentiating into T cells.

The next step will be to produce cells with long-term renewal and engraftment potential in immunocompromised mice, without the challenging overexpression of numerous transgenes. Indeed, these protocols are still very laborious to perform. Moreover the production of 11kb lentivirus remains difficult and 5 to 7 are necessary. (Doulatov et al., 2013; Sugimura et al., 2017; Vo et al., 2018). The transcriptomic comparison between hiPSC-derived HPSCs with cord blood cells showed that CD34^+^CD43^+^ cells clustered with cord blood cells in contrast to CD34^+^CD43^-^ cells. Nevertheless, we observed that genes shown to be essential for grafting potential in animals by G. Daley’s team, such as the HOXA family and LCOR (Sugimura et al., 2017; Vo et al., 2018) were not expressed in CD34^+^CD43^+^. However, only one study has shown so far the generation of long-term multipotent HSPCs supporting hematopoietic reconstitution and self-renewal in vivo, but through early differentiated cells expressing APLNR, thus more likely an endothelial cell undergoing EHT or a newly formed HSPC (Guyonneau-Harmand et al., 2017). D9 CD34^+^CD43^-^ EB cells had a hemogenic-endothelium profile and expressed APLRN, which may give them the potential to engraft in animals. It would be interesting to transplant D9 CD34^+^CD43^-^ EB cells in immunosuppressed animals and follow their differentiation by immunoprofiling and scRNA sequencing.

After establishing robust hematopoietic differentiation, we analyzed induction of lymphopoiesis. After 20/25 days of co-culture on OP9-DLL1, we obtained CD7^+^, CD5^+^, CD4^+^, CD8^+^ cells, which are major markers of T cells. Interestingly the upregulated pathways in hPSC-Td25 and HSC-Td25 were directed by genes involved in T cell differentiation and in the establishment of the TCR and the MHC complex. Furthermore, we have seen that hiPSC-Td25 and HSC-Td25 differentiated cells possessed the same abilities to differentiate as cord blood HSCs under our culture conditions. Even if the signature of the cells after 25 days of co-culture on OP9-DLL1 was clearly typical of thymic progenitors at the DN3a stage, our culture conditions did not allow at that stage the acquisition of a functional TCR either in HSC-TD25 or in T-iPSC-Td25. As other, we observed in our differentiation some CD56^+^ cells (Flippe et al., 2020; Nishimura et al., 2013; Themeli et al., 2013). One hypothesis could be that the CD56^+^ cells that we observe are part of a niche not supporting T cell development (Nishimura et al., 2013; Themeli et al., 2013). Moreover, studies have shown that high expression of IL-7 could also prevent T cell selection (Awong et al., 2009; De Smedt et al., 2004; Six et al., 2011; Zakrzewski et al., 2006). It would be interesting to eliminate CD56^+^ cells from the culture, decrease the concentration of IL-7 and analyze the expression of the TCR.

Advances in immune cell engineering are opening-up new perspectives for the generation of custom synthetic T cells from hiPSCs. TCR acquisition during differentiation will no longer be an issue since antigenic specificity could be brought by other technical means, such as a transgenic TCR (Minagawa et al., 2018) or the use of chimeric antigen receptor (CARs) providing hiPSC-derived T cells with customizable antigen specificity in an HLA-independent manner. Indeed, Themeli et al showed in 2013 the possibility to insert a CAR into hiPSCs under control of an inducible promoter. The induction of the CAR during the differentiation resulted in the generation of functional CAR-T cells in vitro and in vivo efficient in a cancer model (Themeli et al., 2013). Today, most studies on T cell differentiation from PSCs are done to generate cytotoxic effector T cells in the context of cancer immunotherapy. Indeed, PSCs represent an unlimited source of cells and allow the generation of custom modified T cells. Similarly, the use of Tregs derived from PSCs is also of great interest in the treatment of GVHD or autoimmune diseases with the same advantages (Amini et al., 2020). It has been shown that transduction of FOXP3 and a transgenic TCR coding sequences into miPSCs and then differentiation of these cells led to the generation of functional Tregs secreting regulatory molecules such as IL-10 and TGF-β and enabled control of autoimmunity in an arthritic mouse model (M. Haque et al., 2016; R. Haque et al., 2012).

In conclusion, we generated a protocol for the differentiation of DN3a thymic progenitors from HSC, hESC and T-hiPSCs. Generation of T cell progenitors *in vitro* offers the opportunity to better study and understand lymphopoiesis. since it remains extremely difficult to study the genesis of T cells in humans. Thus, *in vitro* generation of T cells would allow unprecedented investigation of lymphopoiesis to functionally test hypotheses. Altogether, our research represents an excellent model to study the establishment of TCR expression during T cell differentiation in humans. This also opens new opportunities for drug screening, disease modeling and could provide an unlimited number of cells for regenerative medicine and cell therapy.

## MATERIAL & METHODS

### hPSCs lines

7 hiPSCs lines were used in this study, all of them derived from healthy donors, T04.01A, T04.01B, T05.003 and T05.006 came from human male adult T-cells, MiPS209 and MiPS220 came from human female adult dermal fibroblasts (Kilens et al., 2018) and LON71 came from human male adult dermal fibroblasts (Gaignerie et al., 2018). hES H9 (WA09 Lot WB0090) were obtained from the WiCell Research Institute, under authorization RE13-004 from the French embryo research oversight committee, Agence de la Biomédecine were also used in this study. Human somatic cells were obtained after informed consent of patients.

### Tissue culture

hPSCs in feeder-free conditions were cultured on Matrigel (BD/Corning) in mTeSR1 (StemCell Technologies); cells were non-enzymatically dissociated with StemMACS passaging solution XF (Miltenyi Biotec) for passaging.

OP9-DLL1/DLL4 stromal cells were kindly provided by Juan-Carlos Zuniga-Pflucker, University of Toronto. OP9-DLL1/DLL4 cells were cultured in OP9 medium (α-MEM with 20% FBS, 2 mM l-glutamine) on gelatin coated plates.

All cells were cultured at 37°C under 20%O2, 5% and were tested for mycoplasma contamination regularly.

### Reprogramming of human T-cells into T-iPSCs

Human T-iPSCs were established from T-cells. In brief, hPBMCs were stimulated with coated anti-CD3 and soluble anti-CD28 MAbs (1 µg/ml each) in complete T-cell medium (RPMI1640 medium, 10% AB serum, 1% of non-essential amino acids, 1% of sodium pyruvate, 1% of hepes, 2 mM L-glutamine and 1 % of Penicillin-Streptomycin) supplemented with IL-2 (1,000 U/ml). After four days of stimulation, 98% CD3^+^TCRαβ^+^ cells were obtained in culture. 10^5^ activated T-cell were seeded per well on a 96-well flat bottom plate in complete RPMI1640 medium 10% AB serum, IL-2 (1,000 U/ml) with coated anti-CD3 and soluble anti-CD28 MAbs (1 µg/ml each) and transduced with three Sendai virus vectors encoding for polycistronic Klf4-Oct4-Sox2, cMyc and Klf4 at MOI of 6-6 and 3.6, respectively. At day 3 of transduction, cells were seeded on 35mm dishes coated with irradiated mouse feeder cells and cultured in T-cell medium. The medium was changed to hPSCs medium (DMEM/F12 supplemented with 20% knockout serum replacer, 2 mM L-glutamine, 1% nonessential amino acids, 10 µM 2-mercaptoethanol, and 5 ng/ml basic fibroblast growth factor (FGF2) on day 5 and was refreshed daily. T-iPSC colonies appeared at ∼13-25 days after transduction. All T-iPSC lines were mechanically passaged to fresh feeder-coated tissue culture dishes with hPSC medium every 6-7 days. After about 10 passages, colonies were picked on MEF and put directly on Matrigel-coated dishes in TeSR1 medium. We waited about 5-6 passes before use for differentiation.

### Early germ layer differentiation

hPSC lines were differentiated into endoderm, mesoderm and ectoderm using Stemmacs Trilineage Kit (Miltenyi biotec). 80,000 cells for mesoderm, 130,000 cells for endoderm and 100,000 cells for ectoderm were plated in 24 wells plates, and cultured in specific media for 7 days, as specified by the protocol. On day 7, differentiated cells were lysed and analyzed by ARN-DGE sequencing.

### SNP analysis

DNA was extracted from somatic and iPSCs samples using the QIAGEN QiaAmp kit, according to the manufacturer’s recommendations. The gDNA was quantified and qualified using a nanodrop. 200 ng of gDNA was outsourced to Integragen Company (Evry, France) for karyotype analysis using HumanCore-24-v1 SNP arrays. This array contains over 300,000 probes distributed throughout the genome with a median coverage of one probe every 5700 bases. All genomic positions were based upon Human Genome Build 37 (hg19). DNA samples were hybridized on HumanCore-24-V1 SNP arrays according to the manufacturer’s instructions by Integragen. Analysis was performed with GenomeStudio software. Chromosome abnormalities were determined by visual inspection of logR ratios and B-allele frequencies (BAF) values and comparing parental cells and iPS-derived samples. LogR ratio, the ratio between observed and expected probe intensity, is informative regarding copy number variation (i.e. deletions/duplications) and BAF is informative regarding heterozygosity. We used the SNP data to compute CNV. In particular, this type of chips allows to detect loss of heterozygosity (LOH), an important concern for hiPSC, which is not possible with classical CGH arrays.

### Hematopoietic T cell differentiation from PSCs

For the differentiation of hPSCs to hematopoietic precursors, we used an optimized serum-and feeder-free in vitro differentiation protocol (Flippe, Gaignerie et al, in revision). Briefly, undifferentiated hPSCs colonies were treated with StemMacs passaging solution XF (Miltenyi Biotec) for 3 min, rinse with α-MEM (Life Technologies) and transferred to low-attachment plates to allow for the formation of embryoid bodies (EBs) in mTeSR1(StemCell Technologies). The formation of EBs was facilitated by 18 hours incubation in the presence of 10µM of Y-27632 dihydrochloride (Axon). At day 0, EBs embryoid bodies were cultured in EB medium (StemPro-34, Life Technologies, with 2 mM l-glutamine, 1% nonessential amino acids, 50 µM 2-mercaptoethanol, 1% penicillin-streptomycin and 50 µg/ml ascorbic acid) with 30 ng/ml of hBMP-4. At day 1, EBs were then cultured with hBMP-4 and 5 ng/ml of hFGF2 until day 3 to allow mesoderm induction. Next, hematopoietic specification and expansion was achieved in the presence of hVEGF (20 ng/ml) and a cocktail of hematopoietic cytokines (hSCF 100 ng/ml, hFlt3L 20 ng/ml, hIL-3 20 ng/ml and hFGF2 5 ng/ml) until day 7 and after continued without hFGF2. Day 9 EBs containing hematopoietic progenitor cells were dissociated by treatment with Accutase for 15 min and single cells were then seeded on OP9-DLL1/DLL4 monolayers to allow for their T-lymphoid differentiation in OP9 medium (α-MEM with 20% FBS, 2 mM l-glutamine, 1% nonessential amino acids, 50 µM 2-mercaptoethanol, 1% penicillin-streptomycin and 50 µg/ml ascorbic acid) supplemented with hSCF 10 ng/ml, hIL-7 5 ng/ml and hFlt3L 10 ng/ml. All recombinant factors were purchased from PeproTech with the exception of hBMP-4 from R&D Systems.

### CFU Assay

MethoCult colony-forming unit (CFU) assays were performed with MethoCult SF H4636 (STEM-CELL Technologies) according to the manufacturer’s protocol. Briefly, cells from EBs day 9 were dissociated with accutase and CD34^+^ cells were MACS purified with the human CD34 UltraPure MicroBead Kit (MiltenyiBiotech). After washing, 10 000 cells were resuspended in 300mL EB medium, gently mixed with 3 mL MethoCult and plated into 2 35mm dishes. Cord blood CD34^+^ cells (2C-101, LONZA) were thawed according to the manufacturer’s instructions, washed in DPBS, and 2000 cells were seeded as described above. Cells were incubated at 37°C and 5% CO2 for 14 days. Total colony numbers were quantified in duplicate plates using an inverted light microscope and the average number of CFU per plate was determined. The methocult was then rehydrated to recover the cells. The cells were washed 3x with DPBS and then analyzed by flow cytometry for the expression of CD15, CD14 and CD235a on the different cell types.

### FACS analysis

The following conjugated antibodies were used for flow cytometric phenotyping and analysis: anti-CD34, anti-CD43, anti-KDR, anti-CD45, anti-CD7, anti-CD5, anti-CD4, anti-CD8, anti-CD3, anti-TCRαβ, anti-CD56, anti-CD15, anti-CD14 and anti-CD235a purchased from BD Biosciences. All antibodies were used in a 1:30 dilution. Dead cells were excluded from analysis in all experiments by staining with DAPI. Flow cytometry analysis was done on a LSRII cytometer (BD Biosciences), a FACS CANTO (BD Biosciences) or a FACS CELESTA (BD Biosciences) and analyzed on FlowJo software.

### Extraction of RNA

Total RNA was extracted using RNeasy® columns and DNAse-treated using RNase-free DNase (Qiagen).

### 3’ Digital Gene Expression (3’ DGE) RNA sequencing

Total RNA molecules were extracted from cells with RNeasy-Mini Kits (Qiagen). Protocol of 3’ DGE RNA sequencing was performed as previously described in (Picarda et al. 2017). Libraries were then sequenced on a NovaSeq 6000 (Illumina). Reads 1 encodes for well specific barcodes and Unique Molecular Identifiers (UMI) and Reads 2 encodes for 3’RNA region. Data were aligned along the human genome reference (hg19) and a count matrix was generated by counting sample specific UMI associated with genes for each sample. Samples with less than 200 000 UMI and less than 5000 genes expressed were excluded of the analysis. Then, a batch correction between samples of different experiments was applied. A Principal Component Analysis (PCA) was performed in order to visualized samples repartition by reducing the number of dimensions. Correlation between samples were assessed with Pearson’s linear correlation heatmaps. Higher correlations are marked in yellow and lower correlations are in red. Differentially expressed genes between conditions were calculated using R package Deseq2 (Bioconductor) by first applying a variance stabilizing transformation (vst). Genes with ajusted p-value inferior to 0.05 were considered as differentially expressed genes. Gene expression were visualized with heatmaps that were generated by scaling and center genes expression. Finally, pathways analysis was performed: R package “Fgsea” and databases such as Kegg, Reactome and Gene Ontology were used to identify significantly enriched or depleted groups of genes in each condition. The accession number for DGE-RNA sequencing raw data is n° PRJEB46691.

## ACKNOWLEGMENT

We thank the iPSC core facility of University of Nantes for its support for tissue culture. We thank the core facilities GenoBIRD for performing the DNA sequencing. We also thank Juan-Carlos Zuniga-Pflucker and Mahmood Mohtashami for kindly providing the OP9-DLL1 and DLL4 cells.

## AUTHOR CONTRIBUTIONS

CG and LD: supervision, and project administration. CG, IA and LD: conceptualization and funding acquisition. LF, CG, and LD: methodology and writing – review and editing. LF, CG, and LD: validation. LF and CS: formal analysis. LF and AG: investigation. OB and XS: contributing essential reagent. LF: writing – original draft. All authors contributed to the article and approved the submitted version.

## FUNDING

This work was funded by: the Labex IGO (project «Investissements d’Avenir», ANR-11-LABX-0016-01). This work was supported by “Paris Scientifique région Pays de la Loire” and by “Fondation pour la recherche médicale (FRM)”.

---

**Supplementary Figure 1:**
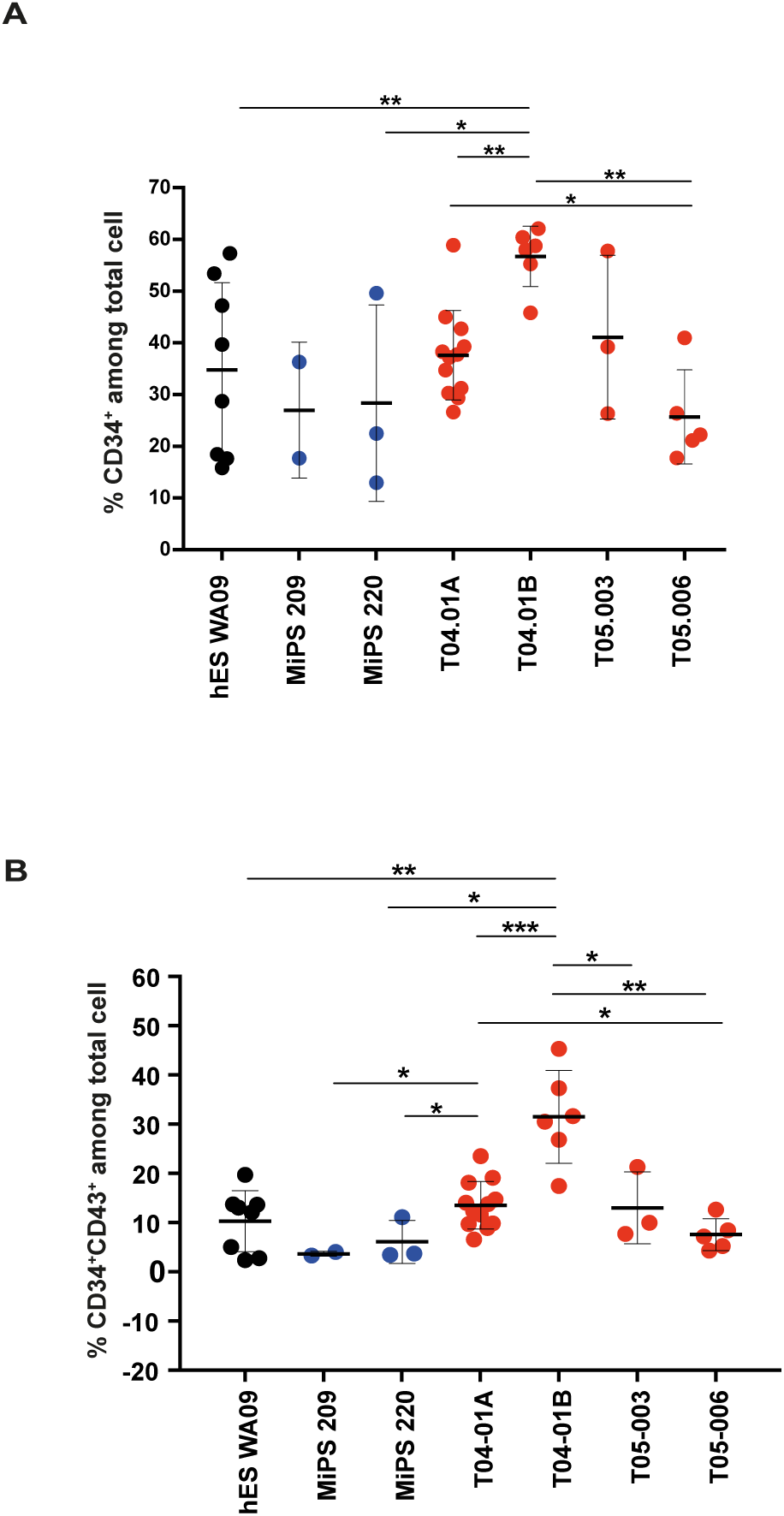
Expression of CD34 and CD43 in HPSCs differentiated from several hiPSCs sources. **A**, Percent of total cells from day 9 EBs expressing CD34, mean +/-SEM are represented. Mann Whitney test, *p<0,05, ***p<0,001, ****p<0, 0001. **B**, Percent of total cells from day 9 EBs expressing CD34 and CD43, mean +/-SEM are represented. Mann Whitney test, *p<0,05, ***p<0,001, ****p<0, 0001.

**Supplementary Figure 2:**
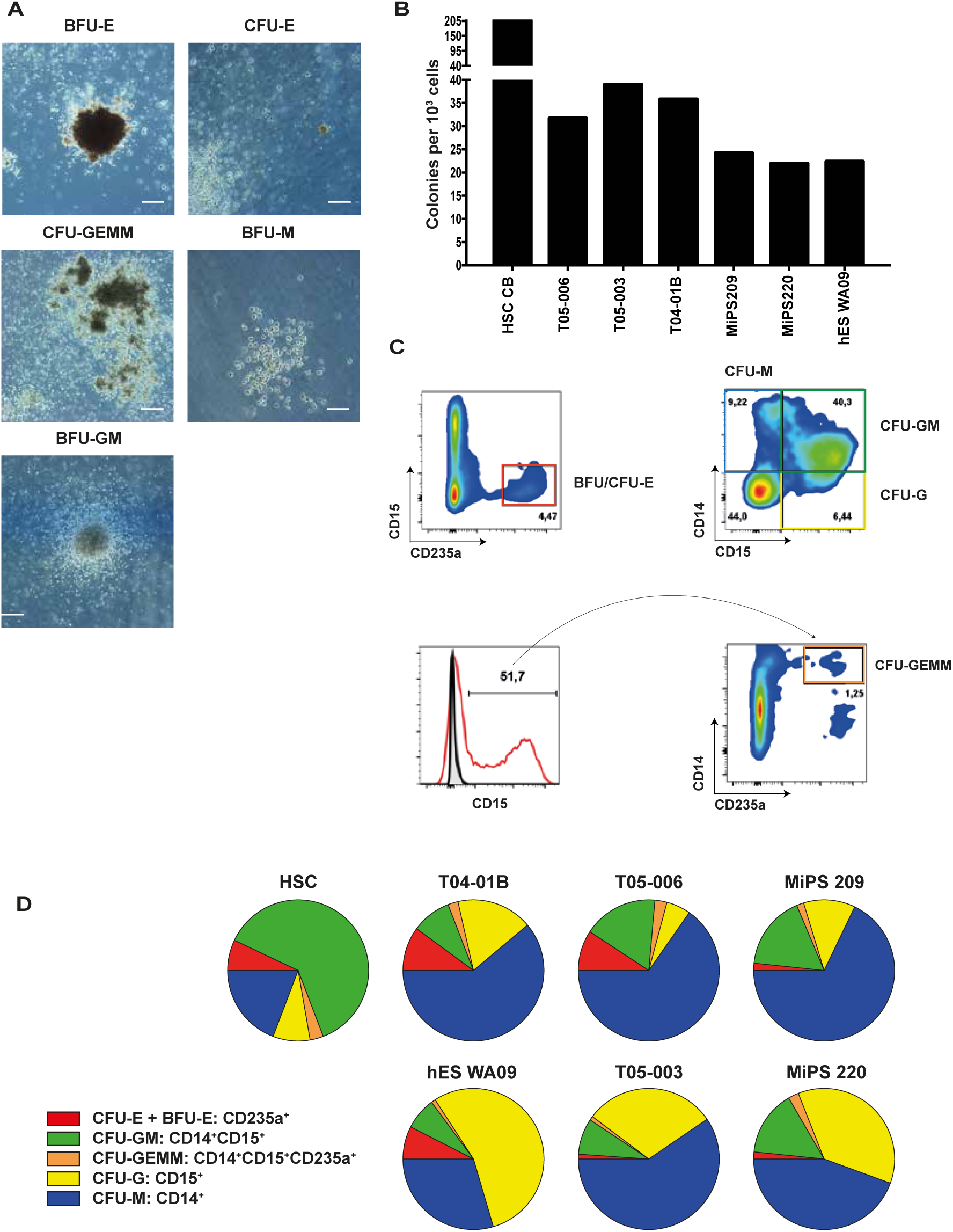
Multipotent HPSCs derived from hPSCs. **A**, Representative photos of blood colonies BFU-E, CFU-E, CFU-GEMM, CFU-M and CFU-GM form hPSCs. **B**, Total colonies number per 1000 CD34+ cells seeded cells in methyl-cellulose in indicated cell lines. **C**, After 14 days in methyl-cellulose, the cells are analyzed by flow-cytometry following the expression of CD14, CD15 and CD235a markers and according to the following gating strategy. The combination of these markers allows to determine the different blood colonies. **D**, Summary of flow-cytometry analysis of CFU assay blood colonies for the indicated cell lines.

**Supplementary Figure 3:**
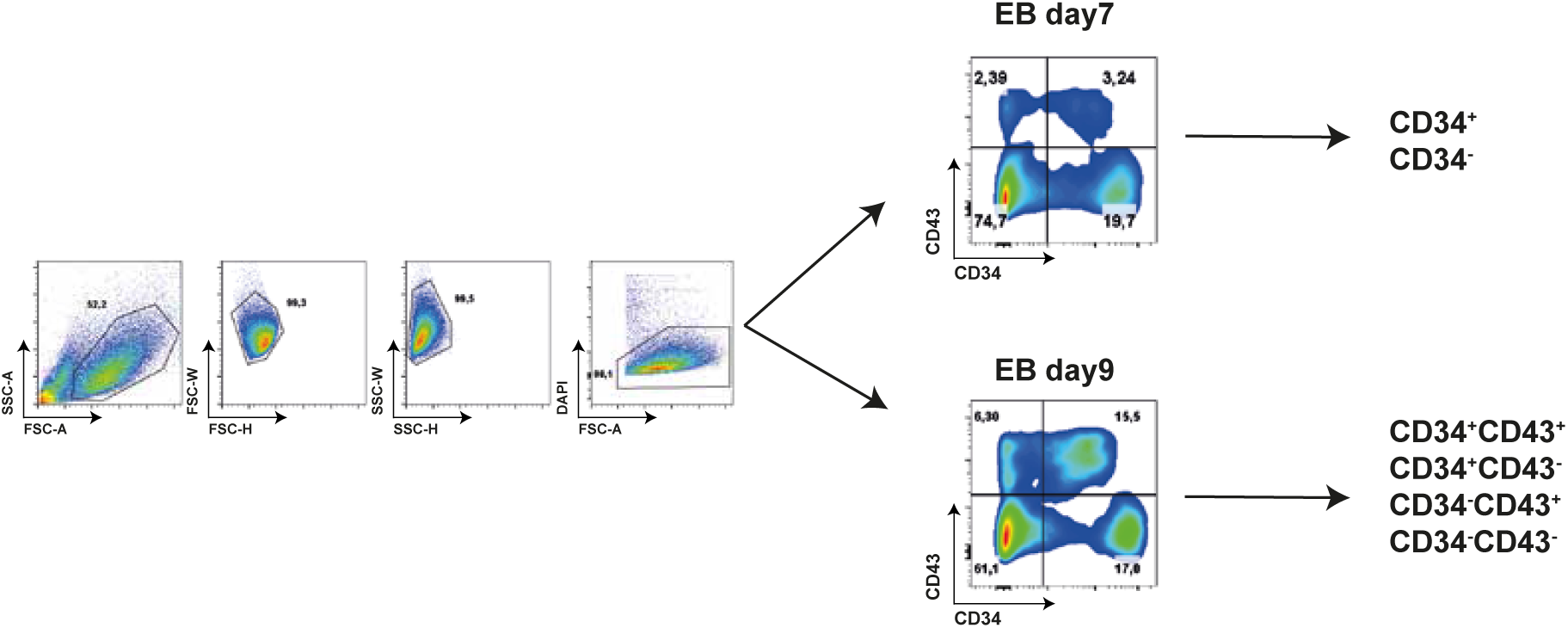
Gating strategy of multipotent HPSCs derived from hPSCs sorting. Gating strategy for cell sorting by FACS Aria. Cells were selected on morphology, exclusion of doublets and dead cells (DAPI), and expression of CD34 et CD43.

**Supplementary Figure 4:**
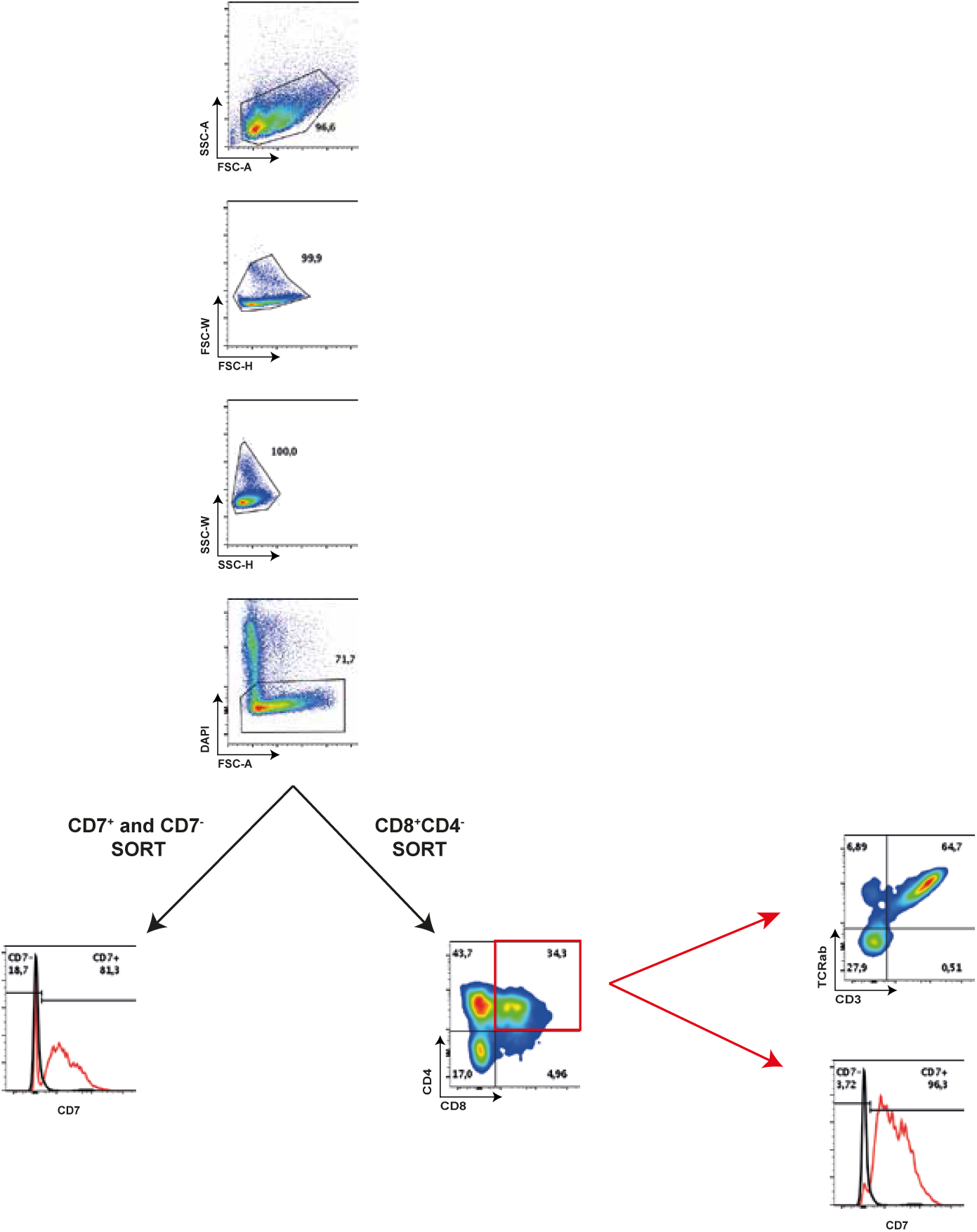
Gating strategy of thymocytes sorting. Gating strategy for cell sorting by FACS Aria. Cells were selected on morphology, exclusion of doublets and dead cells (DAPI), and expression of CD7 or CD8 et CD4. 96,3% of CD8^+^CD4^+^ expressed CD7 and 64,7% CD3 and TCRαβ.

## Notes

### Competing Interest Statement

The authors have declared no competing interest.

## REFERENCES

Amini, L., Greig, J., Schmueck-Henneresse, M., Volk, H.-D., Bézie, S., Reinke, P., Guillonneau, C., Wagner, D. L., & Anegon, I. (2020). Super-Treg: Toward a New Era of Adoptive Treg Therapy Enabled by Genetic Modifications. Frontiers in Immunology, 11, 611638. https://doi.org/10.3389/fimmu.2020.611638

Awong, G., Herer, E., La Motte-Mohs, R. N., & Zúñiga-Pflücker, J. C. (2011). Human CD8 T cells generated in vitro from hematopoietic stem cells are functionally mature. BMC Immunology, 12, 22. https://doi.org/10.1186/1471-2172-12-22

Awong, G., Herer, E., Surh, C. D., Dick, J. E., La Motte-Mohs, R. N., & Zúñiga-Pflücker, J. C. (2009). Characterization in vitro and engraftment potential in vivo of human progenitor T cells generated from hematopoietic stem cells. Blood, 114(5), 972–982. https://doi.org/10.1182/blood-2008-10-187013

Bézie, S., Charreau, B., Vimond, N., Lasselin, J., Gérard, N., Nerrière-Daguin, V., Bellier-Waast, F., Duteille, F., Anegon, I., & Guillonneau, C. (2019a). Human CD8+ Tregs expressing a MHC-specific CAR display enhanced suppression of human skin rejection and GVHD in NSG mice. Blood Advances, 3(22), 3522–3538. https://doi.org/10.1182/bloodadvances.2019000411

Bézie, S., Charreau, B., Vimond, N., Lasselin, J., Gérard, N., Nerrière-Daguin, V., Bellier-Waast, F., Duteille, F., Anegon, I., & Guillonneau, C. (2019b). Human CD8+ Tregs expressing a MHC-specific CAR display enhanced suppression of human skin rejection and GVHD in NSG mice. Blood Advances, 3(22), 3522–3538. https://doi.org/10.1182/bloodadvances.2019000411

Bézie, S., Meistermann, D., Boucault, L., Kilens, S., Zoppi, J., Autrusseau, E., Donnart, A., Nerrière-Daguin, V., Bellier-Waast, F., Charpentier, E., Duteille, F., David, L., Anegon, I., & Guillonneau, C. (2017). Ex Vivo Expanded Human Non-Cytotoxic CD8+CD45RClow/-Tregs Efficiently Delay Skin Graft Rejection and GVHD in Humanized Mice. Frontiers in Immunology, 8, 2014. https://doi.org/10.3389/fimmu.2017.02014

Bissels, U., Aivazidou, F., Knöbel, S., Bosio, A., & Papanikolaou, E. (n.d.). A novel user-independent CFU assay for hematopoietic stem and progenitor cells. 1.

Buono, M., Facchini, R., Matsuoka, S., Thongjuea, S., Waithe, D., Luis, T. C., Giustacchini, A., Besmer, P., Mead, A. J., Jacobsen, S. E. W., & Nerlov, C. (2016). A dynamic niche provides Kit ligand in a stage-specific manner to the earliest thymocyte progenitors. Nature Cell Biology, 18(2), 157–167. https://doi.org/10.1038/ncb3299

Calvanese, V., Nguyen, A. T., Bolan, T. J., Vavilina, A., Su, T., Lee, L. K., Wang, Y., Lay, F. D., Magnusson, M., Crooks, G. M., Kurdistani, S. K., & Mikkola, H. K. A. (2019). MLLT3 governs human haematopoietic stem-cell self-renewal and engraftment. Nature, 576(7786), 281–286. https://doi.org/10.1038/s41586-019-1790-2

Canté-Barrett, K., Mendes, R. D., Li, Y., Vroegindeweij, E., Pike-Overzet, K., Wabeke, T., Langerak, A. W., Pieters, R., Staal, F. J. T., & Meijerink, J. P. P. (2017). Loss of CD44dim Expression from Early Progenitor Cells Marks T-Cell Lineage Commitment in the Human Thymus. Frontiers in Immunology, 8. https://doi.org/10.3389/fimmu.2017.00032

Charpentier, E., Cornec, M., Dumont, S., Meistermann, D., Bordron, P., David, L., Redon, R., Bonnaud, S., & Bihouée, A. (2021). *3’ RNA sequencing for robust and low-cost gene expression profiling* [Preprint]. Protocol Exchange. https://doi.org/10.21203/rs.3.pex-1336/v1

De Smedt, M., Hoebeke, I., & Plum, J. (2004). Human bone marrow CD34+ progenitor cells mature to T cells on OP9-DL1 stromal cell line without thymus microenvironment. Blood Cells, Molecules & Diseases, 33(3), 227–232. https://doi.org/10.1016/j.bcmd.2004.08.007

Demirci, S., Leonard, A., & Tisdale, J. F. (2020). Hematopoietic stem cells from pluripotent stem cells: Clinical potential, challenges, and future perspectives. STEM CELLS Translational Medicine, sctm.20–0247. https://doi.org/10.1002/sctm.20-0247

Desanti, G. E., Jenkinson, W. E., Parnell, S. M., Boudil, A., Gautreau-Rolland, L., Eksteen, B., Ezine, S., Lane, P. J. L., Jenkinson, E. J., & Anderson, G. (2011). Clonal Analysis Reveals Uniformity in the Molecular Profile and Lineage Potential of CCR9 ^+^ and CCR9 ^−^ Thymus-Settling Progenitors. The Journal of Immunology, 186(9), 5227–5235. https://doi.org/10.4049/jimmunol.1002686

Dik, W. A., Pike-Overzet, K., Weerkamp, F., de Ridder, D., de Haas, E. F. E., Baert, M. R. M., van der Spek, P., Koster, E. E. L., Reinders, M. J. T., van Dongen, J. J. M., Langerak, A. W., & Staal, F. J. T. (2005). New insights on human T cell development by quantitative T cell receptor gene rearrangement studies and gene expression profiling. The Journal of Experimental Medicine, 201(11), 1715–1723. https://doi.org/10.1084/jem.20042524

Dong, C., Juedes, A. E., Temann, U. A., Shresta, S., Allison, J. P., Ruddle, N. H., & Flavell, R. A. (2001). ICOS co-stimulatory receptor is essential for T-cell activation and function. Nature, 409(6816), 97–101. https://doi.org/10.1038/35051100

Doulatov, S., Vo, L. T., Chou, S. S., Kim, P. G., Arora, N., Li, H., Hadland, B. K., Bernstein, I. D., Collins, J. J., Zon, L. I., & Daley, G. Q. (2013). Induction of Multipotential Hematopoietic Progenitors from Human Pluripotent Stem Cells via Respecification of Lineage-Restricted Precursors. Cell Stem Cell, 13(4), 459–470. https://doi.org/10.1016/j.stem.2013.09.002

Flippe, L., Bézie, S., Anegon, I., & Guillonneau, C. (2019). Future prospects for CD8+ regulatory T cells in immune tolerance. Immunological Reviews, 292(1), 209–224. https://doi.org/10.1111/imr.12812

Flippe, L., Gaignerie, A., Sérazin, C., Baron, O., Saulquin, X., Themeli, M., Guillonneau, C., & David, L. (2020). Rapid and Reproducible Differentiation of Hematopoietic and T Cell Progenitors From Pluripotent Stem Cells. Frontiers in Cell and Developmental Biology, 8, 577464. https://doi.org/10.3389/fcell.2020.577464

Gaignerie, A., Lefort, N., Rousselle, M., Forest-Choquet, V., Flippe, L., Francois-Campion, V., Girardeau, A., Caillaud, A., Chariau, C., Francheteau, Q., Derevier, A., Chaubron, F., Knöbel, S., Gaborit, N., Si-Tayeb, K., & David, L. (2018). Urine-derived cells provide a readily accessible cell type for feeder-free mRNA reprogramming. Scientific Reports, 8(1), 14363. https://doi.org/10.1038/s41598-018-32645-2

Garcia-Perez, L., Famili, F., Cordes, M., Brugman, M., Eggermond, M. van, Wu, H., Chouaref, J., Granado, D. S. L., Tiemessen, M. M., Pike-Overzet, K., Daxinger, L., & Staal, F. J. T. (2020). Functional definition of a transcription factor hierarchy regulating T cell lineage commitment. Science Advances, 6(31), eaaw7313. https://doi.org/10.1126/sciadv.aaw7313

Georgopoulos, K. (2017). The making of a lymphocyte: The choice among disparate cell fates and the IKAROS enigma. Genes & Development, 31(5), 439–450. https://doi.org/10.1101/gad.297002.117

Georgopoulos, K., Winandy, S., & Avitahl, N. (1997). The role of the Ikaros gene in lymphocyte development and homeostasis. Annual Review of Immunology, 15, 155–176. https://doi.org/10.1146/annurev.immunol.15.1.155

Guyonneau-Harmand, L., L’Homme, B., Birebent, B., Desterke, C., Chevallier, N., Garçon, L., Lapillonne, H., Benderitter, M., Delhommeau, F., Jaffredo, T., Chapel, A., & Douay, L. (2017). Transgene-free hematopoietic stem and progenitor cells from human induced pluripotent stem cells. BioRxiv, 177691. https://doi.org/10.1101/177691

Haque, M., Lei, F., Xiong, X., Das, J. K., Ren, X., Fang, D., Salek-Ardakani, S., Yang, J.-M., & Song, J. (2019). Stem cell-derived tissue-associated regulatory T cells suppress the activity of pathogenic cells in autoimmune diabetes. JCI Insight, 4(7). https://doi.org/10.1172/jci.insight.126471

Haque, M., Song, J., Fino, K., Sandhu, P., Song, X., Lei, F., Zheng, S., Ni, B., Fang, D., & Song, J. (2016). Stem cell-derived tissue-associated regulatory T cells ameliorate the development of autoimmunity. Scientific Reports, 6, 20588. https://doi.org/10.1038/srep20588

Haque, R., Lei, F., Xiong, X., Bian, Y., Zhao, B., Wu, Y., & Song, J. (2012). Programming of regulatory T cells from pluripotent stem cells and prevention of autoimmunity. Journal of Immunology (Baltimore, Md.: 1950), 189(3), 1228–1236. https://doi.org/10.4049/jimmunol.1200633

Kawamoto, H., Masuda, K., Nagano, S., & Maeda, T. (2018a). Cloning and expansion of antigen-specific T cells using iPS cell technology: Development of “off-the-shelf” T cells for the use in allogeneic transfusion settings. International Journal of Hematology, 107(3), 271– 277. https://doi.org/10.1007/s12185-018-2399-1

Kawamoto, H., Masuda, K., Nagano, S., & Maeda, T. (2018b). Cloning and expansion of antigen-specific T cells using iPS cell technology: Development of “off-the-shelf” T cells for the use in allogeneic transfusion settings. International Journal of Hematology, 107(3), 271– 277. https://doi.org/10.1007/s12185-018-2399-1

Kilens, S., Meistermann, D., Moreno, D., Chariau, C., Gaignerie, A., Reignier, A., Lelièvre, Y., Casanova, M., Vallot, C., Nedellec, S., Flippe, L., Firmin, J., Song, J., Charpentier, E., Lammers, J., Donnart, A., Marec, N., Deb, W., Bihouée, A., … Milieu Intérieur Consortium. (2018). Parallel derivation of isogenic human primed and naive induced pluripotent stem cells. Nature Communications, 9(1), 360. https://doi.org/10.1038/s41467-017-02107-w

La Motte-Mohs, R. N., Herer, E., & Zúñiga-Pflücker, J. C. (2005). Induction of T-cell development from human cord blood hematopoietic stem cells by Delta-like 1 in vitro. Blood, 105(4), 1431–1439. https://doi.org/10.1182/blood-2004-04-1293

Lochem, E. G. van, Velden, V. H. J. van der, Wind, H. K., Marvelde, J. G. te, Westerdaal, N. a. C., & Dongen, J. J. M. van. (2004). Immunophenotypic differentiation patterns of normal hematopoiesis in human bone marrow: Reference patterns for age-related changes and disease-induced shifts. Cytometry Part B: Clinical Cytometry, 60B(1), 1–13. https://doi.org/10.1002/cyto.b.20008

Maeda, T., Nagano, S., Ichise, H., Kataoka, K., Yamada, D., Ogawa, S., Koseki, H., Kitawaki, T., Kadowaki, N., Takaori-Kondo, A., Masuda, K., & Kawamoto, H. (2016). Regeneration of CD8αβ T Cells from T-cell-Derived iPSC Imparts Potent Tumor Antigen-Specific Cytotoxicity. Cancer Research, 76(23), 6839–6850. https://doi.org/10.1158/0008-5472.CAN-16-1149

Minagawa, A., Yoshikawa, T., Yasukawa, M., Hotta, A., Kunitomo, M., Iriguchi, S., Takiguchi, M., Kassai, Y., Imai, E., Yasui, Y., Kawai, Y., Zhang, R., Uemura, Y., Miyoshi, H., Nakanishi, M., Watanabe, A., Hayashi, A., Kawana, K., Fujii, T., … Kaneko, S. (2018). Enhancing T Cell Receptor Stability in Rejuvenated iPSC-Derived T Cells Improves Their Use in Cancer Immunotherapy. Cell Stem Cell, 23(6), 850–858.e4. https://doi.org/10.1016/j.stem.2018.10.005

Nianias, A., & Themeli, M. (2019). Induced Pluripotent Stem Cell (iPSC)–Derived Lymphocytes for Adoptive Cell Immunotherapy: Recent Advances and Challenges. Current Hematologic Malignancy Reports, 14(4), 261–268. https://doi.org/10.1007/s11899-019-00528-6

Nishimura, T., Kaneko, S., Kawana-Tachikawa, A., Tajima, Y., Goto, H., Zhu, D., Nakayama-Hosoya, K., Iriguchi, S., Uemura, Y., Shimizu, T., Takayama, N., Yamada, D., Nishimura, K., Ohtaka, M., Watanabe, N., Takahashi, S., Iwamoto, A., Koseki, H., Nakanishi, M., … Nakauchi, H. (2013). Generation of rejuvenated antigen-specific T cells by reprogramming to pluripotency and redifferentiation. Cell Stem Cell, 12(1), 114–126. https://doi.org/10.1016/j.stem.2012.11.002

Outters, P., Jaeger, S., Zaarour, N., & Ferrier, P. (2015). Long-Range Control of V(D)J Recombination & Allelic Exclusion: Modeling Views. Advances in Immunology, 128, 363– 413. https://doi.org/10.1016/bs.ai.2015.08.002

Paes, B. C. M. F., Moço, P. D., Pereira, C. G., Porto, G. S., de Sousa Russo, E. M., Reis, L. C. J., Covas, D. T., & Picanço-Castro, V. (2017). Ten years of iPSC: Clinical potential and advances in vitro hematopoietic differentiation. Cell Biology and Toxicology, 33(3), 233– 250. https://doi.org/10.1007/s10565-016-9377-2

Park, J.-E., Botting, R. A., Domínguez Conde, C., Popescu, D.-M., Lavaert, M., Kunz, D. J., Goh, I., Stephenson, E., Ragazzini, R., Tuck, E., Wilbrey-Clark, A., Roberts, K., Kedlian, V. R., Ferdinand, J. R., He, X., Webb, S., Maunder, D., Vandamme, N., Mahbubani, K. T., … Teichmann, S. A. (2020). A cell atlas of human thymic development defines T cell repertoire formation. Science (New York, N.Y.), 367(6480). https://doi.org/10.1126/science.aay3224

Plath, K., & Lowry, W. E. (2011). Progress in understanding reprogramming to the induced pluripotent state. Nature Reviews. Genetics, 12(4), 253–265. https://doi.org/10.1038/nrg2955

Rothenberg, E. V., Kueh, H. Y., Yui, M. A., & Zhang, J. A. (2016). Hematopoiesis and T-cell specification as a model developmental system. Immunological Reviews, 271(1), 72–97. https://doi.org/10.1111/imr.12417

Rothenberg, E. V., Moore, J. E., & Yui, M. A. (2008). Launching the T-cell-lineage developmental programme. Nature Reviews. Immunology, 8(1), 9–21. https://doi.org/10.1038/nri2232

Schmitt, T. M., & Zúñiga-Pflücker, J. C. (2002). Induction of T cell development from hematopoietic progenitor cells by delta-like-1 in vitro. Immunity, 17(6), 749–756.

Six, E. M., Benjelloun, F., Garrigue, A., Bonhomme, D., Morillon, E., Rouiller, J., Cacavelli, L., Blondeau, J., Beldjord, K., Hacein-Bey-Abina, S., Cavazzana-Calvo, M., & André-Schmutz, I. (2011). Cytokines and culture medium have a major impact on human in vitro T-cell differentiation. Blood Cells, Molecules & Diseases, 47(1), 72–78. https://doi.org/10.1016/j.bcmd.2011.04.001

Sugimura, R., Jha, D. K., Han, A., Soria-Valles, C., da Rocha, E. L., Lu, Y.-F., Goettel, J. A., Serrao, E., Rowe, R. G., Malleshaiah, M., Wong, I., Sousa, P., Zhu, T. N., Ditadi, A., Keller, G., Engelman, A. N., Snapper, S. B., Doulatov, S., & Daley, G. Q. (2017). Haematopoietic stem and progenitor cells from human pluripotent stem cells. Nature, 545(7655), 432–438. https://doi.org/10.1038/nature22370

Themeli, M., Kloss, C. C., Ciriello, G., Fedorov, V. D., Perna, F., Gonen, M., & Sadelain, M. (2013). Generation of tumor-targeted human T lymphocytes from induced pluripotent stem cells for cancer therapy. Nature Biotechnology, 31(10), 928–933. https://doi.org/10.1038/nbt.2678

Themeli, M., Rivière, I., & Sadelain, M. (2015). New cell sources for T cell engineering and adoptive immunotherapy. Cell Stem Cell, 16(4), 357–366. https://doi.org/10.1016/j.stem.2015.03.011

Vo, L. T., Kinney, M. A., Liu, X., Zhang, Y., Barragan, J., Sousa, P. M., Jha, D. K., Han, A., Cesana, M., Shao, Z., North, T. E., Orkin, S. H., Doulatov, S., Xu, J., & Daley, G. Q. (2018). Regulation of embryonic haematopoietic multipotency by EZH1. Nature, 553(7689), 506– 510. https://doi.org/10.1038/nature25435

Vodyanik, M. A., Thomson, J. A., & Slukvin, I. I. (2006). Leukosialin (CD43) defines hematopoietic progenitors in human embryonic stem cell differentiation cultures. Blood, 108(6), 2095–2105. https://doi.org/10.1182/blood-2006-02-003327

Wang, H., Kadlecek, T. A., Au-Yeung, B. B., Goodfellow, H. E. S., Hsu, L.-Y., Freedman, T. S., & Weiss, A. (2010). ZAP-70: An Essential Kinase in T-cell Signaling. Cold Spring Harbor Perspectives in Biology, 2(5). https://doi.org/10.1101/cshperspect.a002279

Wang, M., Wang, H., Wen, Y., Chen, X., Liu, X., Gao, J., Su, P., Xu, Y., Zhou, W., Shi, L., & Zhou, J. (2018). MEIS2 regulates endothelial to hematopoietic transition of human embryonic stem cells by targeting TAL1. Stem Cell Research & Therapy, 9(1), 340. https://doi.org/10.1186/s13287-018-1074-z

Yui, M. A., & Rothenberg, E. V. (2014). Developmental gene networks: A triathlon on the course to T cell identity. Nature Reviews. Immunology, 14(8), 529–545. https://doi.org/10.1038/nri3702

Zakrzewski, J. L., Kochman, A. A., Lu, S. X., Terwey, T. H., Kim, T. D., Hubbard, V. M., Muriglan, S. J., Suh, D., Smith, O. M., Grubin, J., Patel, N., Chow, A., Cabrera-Perez, J., Radhakrishnan, R., Diab, A., Perales, M.-A., Rizzuto, G., Menet, E., Pamer, E. G., … van den Brink, M. R. M. (2006). Adoptive transfer of T-cell precursors enhances T-cell reconstitution after allogeneic hematopoietic stem cell transplantation. Nature Medicine, 12(9), 1039–1047. https://doi.org/10.1038/nm1463

Zhou, W., Yui, M. A., Williams, B. A., Yun, J., Wold, B. J., Cai, L., & Rothenberg, E. V. (2019). Single-Cell Analysis Reveals Regulatory Gene Expression Dynamics Leading to Lineage Commitment in Early T Cell Development. Cell Systems, 9(4), 321–337.e9. https://doi.org/10.1016/j.cels.2019.09.008

